# Evaluating the Role of Fish as Surrogates for Amphibians in Pesticide Ecological Risk Assessment

**DOI:** 10.1101/584417

**Authors:** Scott Glaberman, Jean Kiwiet, Catherine Aubee

## Abstract

Ecological risk of chemicals to aquatic-phase amphibians has historically been evaluated by comparing estimated environmental concentrations in surface water to surrogate toxicity data from standard fish species. Despite their obvious similarities, there are biological disparities among fish and amphibians that could affect their exposure and response to chemicals. Given the alarming decline in amphibians in which anthropogenic pollutants play at least some role, evaluating the potential risk of chemicals to amphibians is becoming increasingly important. Here, we evaluate relative sensitivity of fish and larval aquatic-phase amphibians to 45 different pesticides using existing data for three standardized toxicity tests: (1) amphibian metamorphosis assay (AMA) with the African clawed frog (*Xenopus laevis*); (2) fish short-term reproductive assay (FSTRA) with freshwater fathead minnow (*Pimephales promelas*); (3) fish early life stage test with *P. promelas* or rainbow trout (*Oncorhynchus mykiss*). The advantage of this dataset over previous work is that these studies show high consistency in exposure method and exposure concentration validation, study duration, test species, endpoints measured, and number of concentrations tested. We found very strong positive relationships between fish and tadpole lowest adverse effect concentrations (LOAEC) for survival (r^2^=0.85, slope=0.97), body weight (r^2^=0.77, slope=0.98), and length (r^2^=0.77, slope=0.92) with only one out of 45 chemicals exhibiting 100-folder greater sensitivity in frogs relative to fish. While these results suggest comparable toxicity for pesticides between these two groups of vertebrates, testing with a greater diversity of amphibians will help determine the generalizability of these results across all amphibians.

**DISCLAIMER:** The views expressed in this manuscript are solely those of the authors and do not represent the policies of the U.S. Environmental Protection Agency. Mention of trade names of commercial products should not be interpreted as an endorsement by the U.S. Environmental Protection Agency.

## INTRODUCTION

Ecological risk of chemicals to aquatic-phase amphibians has historically been evaluated by comparing estimated environmental concentrations in surface water to surrogate toxicity data from standard fish species (Ortiz-Santaliestra *et al*., 2017; U.S. EPA, 2015a). Although the origin and rationale for this common practice is unclear, it is likely based, in part, on the many physical and life history similarities between the two groups including ectothermy and their reliance on aquatic environments for some or all stages of their life cycle including reproduction. In addition, while there have been historical efforts to establish ecotoxicology models in amphibians [*e.g*., frog embryo teratogenesis assay – *Xenopus* (FETAX); (Dumont *et al*., 1983)], there are few if any such models with the breadth of toxicity data and method development that could rival major fish models such as fathead minnows, rainbow trout, or zebrafish (*Danio rerio*). Given the decline in some species of amphibians in which anthropogenic pollutants may play at least some role (Blaustein *et al*., 2011), evaluating the risk of chemicals to amphibians is becoming increasingly important.

Despite their obvious similarities, there are biological disparities among fish and amphibians. First, if evolutionary divergence time is used as a marker for the potential accumulation of biological differences among taxa, then the estimated 450 million year separation between extant ray-finned fish (which includes 99% of all fish species) and amphibians (Irisarri *et al*., 2017), provides substantial time for increased diversity both within and between these major vertebrate groups. For example, many adult amphibians undergo gas and ion exchange through their highly permeable skin and/or lungs, while larval amphibians respire through their skin and/or gills (Boutilier *et al.*, 1992; Boyer and Grue, 1995). Some amphibians no longer have lungs but rather modified body surfaces or altered habitat requirements for improved cutaneous respiration (Wells, 2010). Most fish rely primarily on gills or a venous plexus on their airbladders (Randall, 1982). Since absorption of chemicals across dermal or respiratory surfaces can be an important route of chemical exposure (Willens *et al.*, 2006), this is just one example of how the diversity between these two vertebrate groups could lead to differences in potential exposure and response to chemicals.

Given the growing concern for protecting amphibian diversity and the biological differences between amphibians and fish, attention has been devoted to comparing chemical sensitivity among these two groups (reviewed in Johnson *et al*., 2017; Weltje *et al.*, 2013). Overall, there is little evidence that amphibians are more sensitive than fish, with the exception of some metals (Birge *et al*., 2000; Hoke and Ankley, 2005) and phenolic compounds (Kerby *et al.*, 2010). In a more recent study, Weltje *et al*. (2013) noted that previous amphibian-fish toxicity comparisons focus almost exclusively on acute toxicity data, which may not be reflective of realistic environmental exposure scenarios. Consequently, they paired chronic toxicity data for fish and aquatic-phase amphibians from a range of test species and endpoints for 52 chemicals compiled from a mixture of regulatory and published literature studies accessed through U.S. EPA’s ECOTOXicity Knowledgebase (U.S. EPA, 2018). After vetting endpoint reliability, out of the 52 chemicals considered, there were three instances where amphibians were more sensitive. Amphibians were 10- to 100-fold more sensitive than fish for two chemicals (carbaryl and dexamethasone), and greater than 100-fold more sensitive for one chemical (sodium perchlorate). Therefore, given the few studies where amphibians were more sensitive, the authors concluded that “additional amphibian testing is not necessary during chemical risk assessment (Weltje *et al*., 2013).”

One of the major challenges in comparing the sensitivities of amphibian and fish to chemicals has been the lack of consistent toxicity data in terms species and life stage tested, experimental design, endpoint choice, and the use of nominal versus measured exposure concentrations. This is primarily because fish toxicity studies are underpinned by longstanding regulatory requirements (U.S. EPA, 2017a) and study guidelines (OECD, 2019; U.S. EPA, 2017b), while standardized protocols for amphibian testing have lagged behind. For example, Weltje *et al*. (2013) analyzed aquatic-phase amphibian chronic toxicity data from a pool of 14 species that included a mixture of apical endpoints (survival, growth, reproduction, development) and life stages with exposure durations ranging from 10 to 210 days. While casting such a wide net is practical and arguably necessary given the relative dearth of amphibian toxicity data, such studies may not be comparable across chemicals for all the reasons mentioned above.

Recently, the U.S. EPA requested screening data for an initial battery of 45 chemicals for both the fish short-term reproductive assay (FSTRA) and the amphibian metamorphosis assay (AMA) through Endocrine Disruptor Screening Program (U.S. EPA, 2009a) mandated through the Food Quality Protection Act. The AMA (OECD, 2009; U.S. EPA, 2009b) is conducted with *X. laevis* starting at the tadpole stage while the FSTRA (OECD, 2012; U.S. EPA, 2009c) is conducted with *P. promelas* beginning when the fish become reproductively active. Both tests have 21-day chemical exposure durations and both include apical measures including survival and growth, in addition to a suite of endocrine-specific endpoints. Although these studies were designed as screening studies (EDSP Tier 1) for detecting endocrine disruption within a weight-of-evidence framework, and not necessarily to evaluate concentration-response given the limited number of concentrations tested and the spacing of test concentrations, the consistency in which the data are collected across chemicals on the same species over the same study duration lends critical comparability for these data that has been lacking in previous fish-amphibian evaluations. Here, we reanalyze these data, focusing on apical endpoints (growth and survival) to evaluate the relative sensitivity of fish and amphibian models and possible implications for ecological risk assessment. We also compare AMA toxicity data to the fish early life stage (ELS) study (OECD, 2013; U.S. EPA, 2016), which is conducted with newly fertilized embryos of *P. promelas* or *O. mykiss* to the juvenile life stage. The ELS provides important grounding for our analysis since the study 1) has served as the common fish chronic toxicity test over the last several decades, 2) offers a second fish dataset with which to compare to the FSTRA, and 3) yields data from an earlier fish life stage as compared to the FSTRA.

## MATERIALS AND METHODS

### Data Acquisition

We extracted endpoints for survival, body weight, and length (total length for fish; snout-to-vent length for tadpoles) from publicly available U.S. EPA reviews of chemical industry data submissions for 45 different pesticide technical grade active ingredients (U.S. EPA, 2015b). We considered data generated according to amphibian metamorphosis assay (AMA), fish short-term reproduction assay (FSTRA), and fish early life stage (ELS) toxicity test designs (Table 1). Each laboratory study included at least 21 days of chemical exposure and was conducted according to standard U.S. EPA test guidelines (U.S. EPA, 2009b, 2009c, 2016). Each study underwent at least three rounds of U.S EPA scientific review. Mode of action (MOA) was assigned to each chemical based on U.S. EPA’s MOATox classification scheme (Barron *et al*., 2015).

**Table 1.** Comparison of Amphibian and Fish Toxicity Test Designs

|  | **Amphibian**<br>**Metamorphosis Assay**<br>**(AMA)** | **Fish Short Term**<br>**Reproduction Assay**<br>**(FSTRA)** | **Fish Early Life Stage**<br>**Assay**<br>**(ELS)** |
| --- | --- | --- | --- |
| *U.S. EPA*<br>*Guideline* | 890.1100 | 890.1350 | 850.1400 |
| *OECD Guideline* | 231 | 229 | 210 |
| *Species* | *Xenopus laevis* | *Pimephales Promelas* | *Pimephales Promelas*<br>or *Oncorhynchus mykiss* |
| *Life Stage* | Tadpole (NF stage 51) | Reproductively active<br>(4.5-6 months old) | Post-fertilization until<br>early juvenile<br>development |
| *Duration* | 21 days | 21 days | 28-32 days (warm<br>water)<br>70-90 days (cold water) |
| *No. of Test*<br>*Concentrations* | ≥3 | ≥3 | ≥5 |
| *Relevant*<br>*Endpoints* | Survival, snout-vent<br>length, body weight | Survival, total length,<br>body weight | Survival, total, body<br>weight |

### Data Analysis

All comparisons were performed based on the lowest observed adverse effect concentration (LOAEC) for survival, length, and body weight endpoints derived from each study. The LOAEC was identified as the lowest test concentration in which statistically significant (p<0.05) adverse effects were detected. All endpoints were based on measured test concentrations (mean-measured or time-weighted average). In cases where the LOAEC was also the lowest concentration tested or, conversely, there were no statistically significant effects at the highest treatment level, the LOAEC was explicitly labeled as “unbounded” [following the terminology in (Weltje *et al*., 2013)] in all analyses.

Linear models comparing fish toxicity endpoints to corresponding amphibian endpoints for the same chemicals were run in R (R Core Team, 2016). We also used a paired t-test to examine whether amphibians were more sensitive than fish in standard studies (α=0.05). Results were further split by product type (herbicide, fungicide, insecticide). Summary statistics were generated to describe the number of cases in which the amphibian endpoint was lower and for the magnitude of difference between endpoints for a given chemical. Additional linear regression analyses were also conducted between FSTRA and ELS study designs and between lowest and highest concentrations tested in fish versus frog studies as measures of consistency among studies.

## RESULTS

### Dataset Composition and Study Design

Of the 45 pesticides in the overall dataset, there are 22 insecticides, 11 fungicides, 10 herbicides, and 2 inert chemicals (Supplemental Table 1). In terms of MOA, 16 chemicals act via narcosis, 10 via acetylcholinesterase inhibition, 7 via other neurotoxic mechanisms, and 2 via reactivity (Barron *et al.*, 2015).

In general, the concentration range tested between fish and larval frog studies are comparable, with lowest and highest test concentrations often within an order of magnitude between the two groups of organisms (Fig. 1). The concentrations tested are more evenly matched between the FSTRA and AMA studies, which reflect similarities in study design as recommended by test guidelines. Moreover, ELS studies often use a narrower range of test concentrations than AMA studies. Linear regression analysis indicates a strong linear relationship between low (r^2^=0.90; p<0.01; intercept=-0.05; slope=0.98) and high (r^2^=0.91; p<0.01; intercept=0.02; slope=1.0) concentrations tested in the FSTRA versus AMA, while the low (r^2^=0.65; p<0.01; intercept=-0.68; slope=0.87) and high (r^2^=0.75; p<0.01; intercept=0.04; slope=0.98) concentrations tested in the ELS and AMA are also strong, but not as robust. As an indication of concentration spacing, the mean (and median) ratio of highest to lowest log concentrations tested in the FSTRA, AMA, and ELS studies are 1.65 (1.84), 1.66 (1.87), and 1.15 (1.20), respectively, despite the fact that the ELS may contain more test concentrations (generally up to five) with narrower spacing between each treatment level (e.g., EPA 850 guidelines require spacing <3.2x; (U.S. EPA, 2016)).

**Figure 1.**
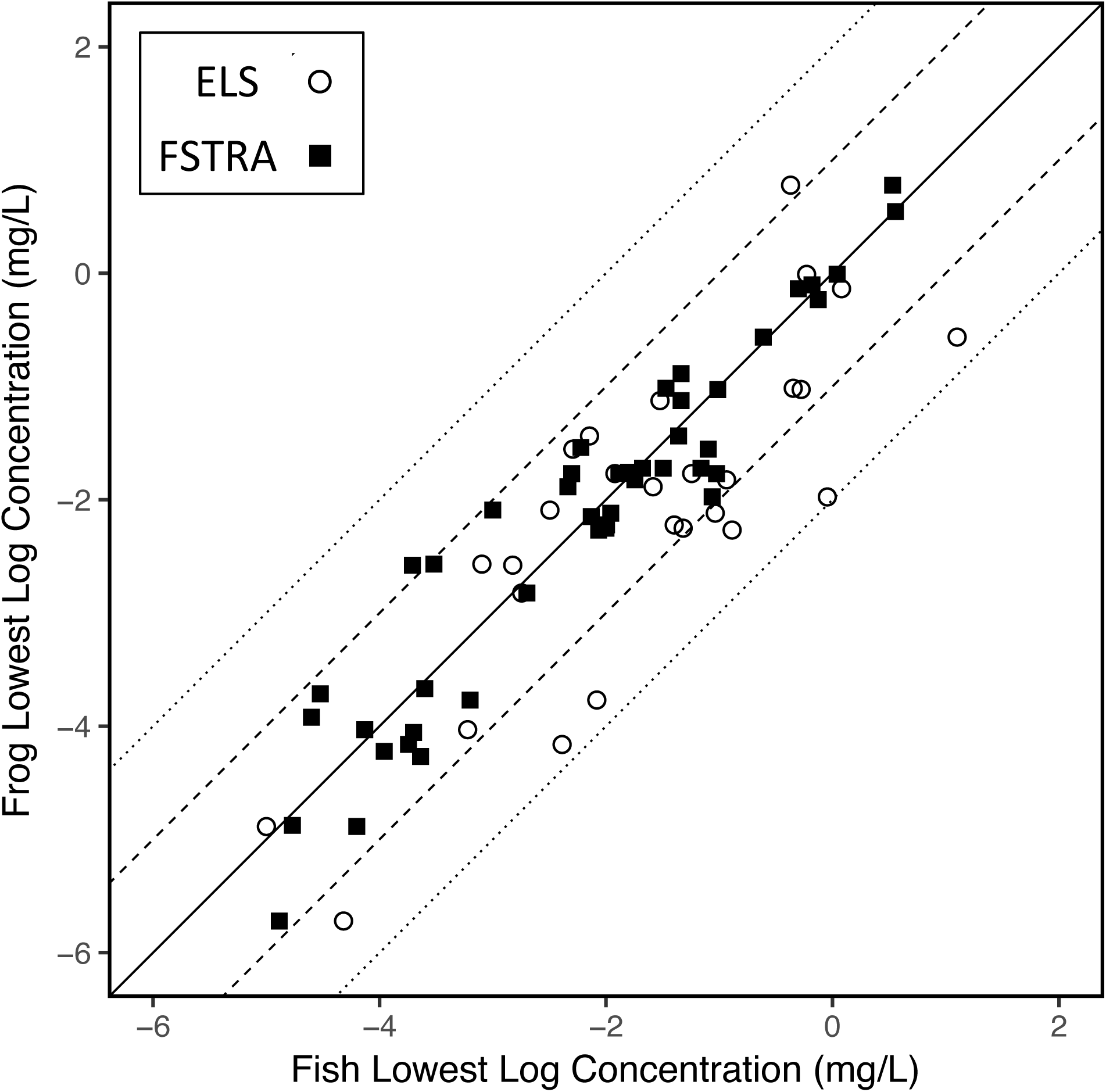
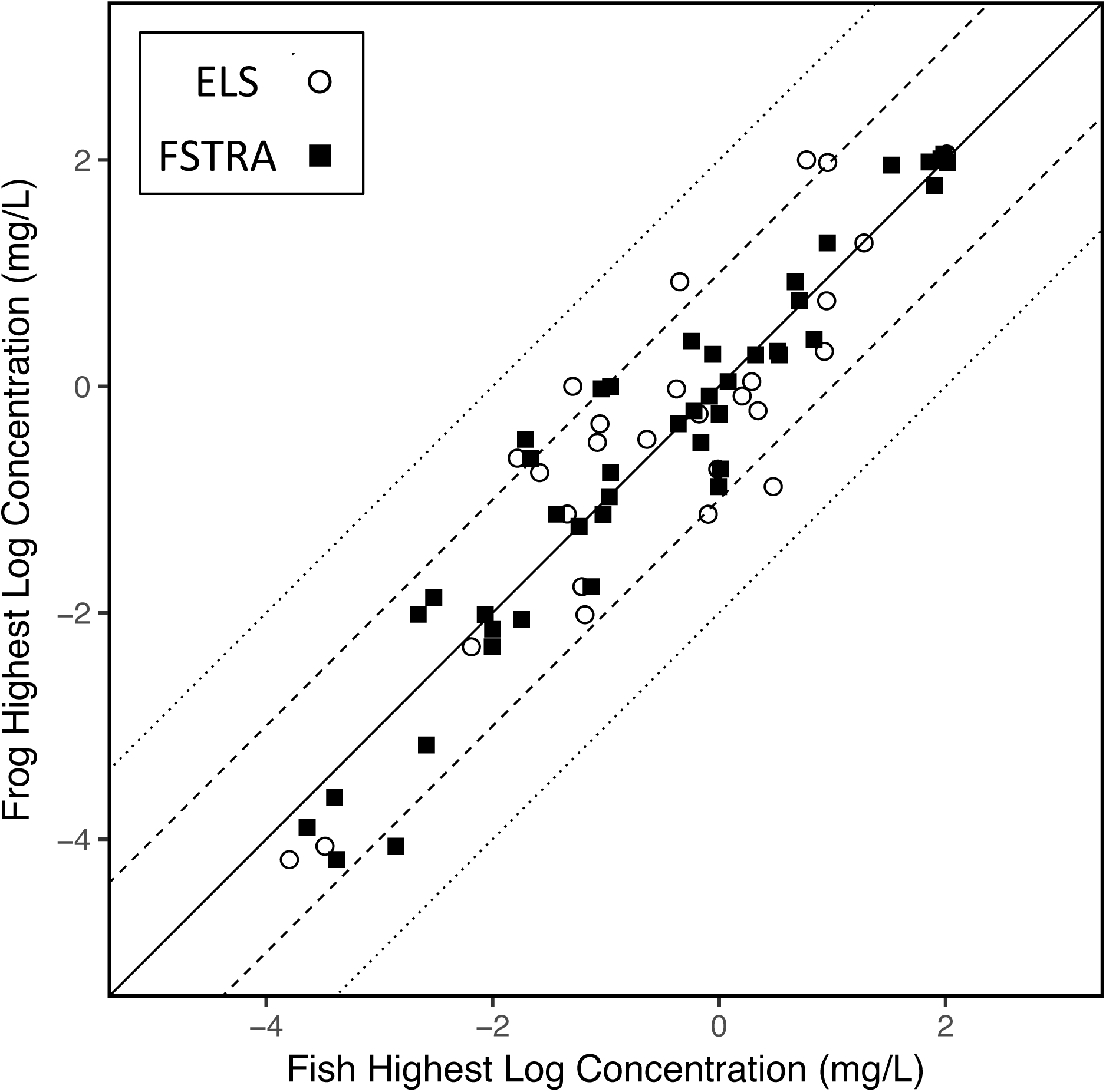
(a) Lowest and (b) highest concentrations tested in fish (FSTRA and ELS) and frog (AMA) toxicity studies. Dashed and dotted lines represent one and two order of magnitude differences, respectively, in concentrations tested between the two groups of organisms.

### Comparison of AMA, FSTRA, and ELS Endpoints

Based on LOAEC comparisons between FSTRA and AMA tests across pesticides, there is a strong linear relationship between fish and tadpoles for body weight, length, and survival, with coefficients of determination (*r*^*2*^) ≥0.77 (Table 2). In comparison, the relationship between the ELS and AMA is also significant, but relatively weak across the three endpoints, with *r*^*2*^ ≥0.51 (Table 2).

**Table 2.** Summary of Pesticide Endpoint Comparisons for Fish Short-Term Reproduction Assay (FSTRA), Amphibian Metamorphosis Assay (AMA), and Fish Early Life Stage Test (ELS).

| Endpoint | Group 1<br>(Predictor) | Group 2<br>(Response) | Linear Regression |  |  | Paired T-Test | Proportion of Chemicals w/ Greater<br>Sensitivity in Tadpole (*X. laevis*) |  |  |
| --- | --- | --- | --- | --- | --- | --- | --- | --- | --- |
| | | | Coeff. of<br>Determ. ($r^2$) | Slope | Intercept | Log Mean of Differences:<br>Group 1 - Group 2<br>(95% CI) | >1x More<br>Sensitive | >10x More<br>Sensitive | >100x More<br>Sensitive |
| Body<br>Weight | FSTRA | AMA | 0.77* | 0.98 | 0.07 | -0.09<br>(-0.34 – 0.16) | 0.49<br>(22 of 45) | 0.09<br>(4 of 45) | 0.00<br>(0 of 45) |
|  | ELS | AMA | 0.54* | 0.81 | 0.04 | -0.20<br>(-0.63 – 0.24) | 0.44<br>(12 of 27) | 0.15<br>(4 of 27) | 0.04<br>(1 of 27) |
|  | FSTRA | ELS | 0.68* | 0.92 | -0.22 | 0.06<br>(-0.26 – 0.38) | -- | -- | -- |
| Length | FSTRA | AMA | 0.77* | 0.92 | -0.13 | 0.08<br>(-0.18 – 0.35) | 0.54<br>(22 of 41) | 0.15<br>(6 of 41) | 0.00<br>(0 of 41) |
|  | ELS | AMA | 0.51* | 0.83 | 0.13 | -0.24<br>(-0.66 – 0.18) | 0.44<br>(11 of 25) | 0.12<br>(3 of 25) | 0.00<br>(0 of 25) |
|  | FSTRA | ELS | 0.77* | 0.80 | -0.41 | 0.33<br>(0.03 – 0.63)* | -- | -- | -- |
| Survival | FSTRA | AMA | 0.85* | 0.97 | 0.02 | -0.03<br>(-0.24 – 0.17) | 0.44<br>(20 of 45) | 0.04<br>(2 of 45) | 0.02<br>(1 of 45) |
|  | ELS | AMA | 0.66* | 0.91 | 0.21 | -0.28<br>(-0.64 – 0.08) | 0.41<br>(11 of 27) | 0.04<br>(1 of 27) | 0.04<br>(1 of 27) |
|  | FSTRA | ELS | 0.83* | 0.92 | -0.35 | 0.31<br>(0.08 – 0.54)* | -- | -- | -- |

\* Statistically significant ( $p < 0.05$ )

Of the 45 pesticides compared, the AMA results in a lower endpoint than the FSTRA approximately half the time (49%, 54%, and 44% of the time for body weight, length, and survival, respectively; Table 2). However, when evaluated statistically, the paired t-test results indicate that there is no significant difference (p>0.05) between FSTRA and AMA endpoint values for the dataset as a whole (Table 2) or when endpoints are further subdivided by pesticide type (not shown). Although not statistically significant, body weight is >10-fold more sensitive in frogs as compared to fish for 4 (9%) out of 45 chemicals; length is >10-fold more sensitive in tadpoles for 6 (15%) out of 41 chemicals; finally, survival is >10-fold more sensitive in tadpoles for 2 (4%) out of 45 chemicals (Table 2). No pesticides are >100-fold more sensitive in tadpoles for body weight or length, while only a single chemical, the fungicide propiconazole, is >100-fold more sensitive in tadpoles for survival. When body weight, length, and survival endpoints are pooled, and the lowest (most sensitive) overall endpoint is used for both FSTRA and AMA studies, the latter is more sensitive 54% of the time and no chemicals are >100-fold more sensitive in the tadpole. Thus, the endpoint-by-endpoint comparisons and the lowest overall endpoint comparison yield concordant results in terms of relative fish-to-frog sensitivity using the FSTRA and AMA.

Although no chemicals in the FSTRA versus AMA comparison were identified as being >100-fold more sensitive in tadpoles based on LOAEC values for growth, several pesticides were >10-fold more sensitive in tadpoles and also associated with unbounded endpoint values that point to potentially underestimated relative sensitivity in tadpoles. This includes cases in which significant effects are observed at the highest test concentration in tadpoles but not fish, and/or 2) significant effects are observed at the lowest test concentration in the tadpoles but not fish. Both of these scenarios could result in a potentially greater relative sensitivity in tadpoles. Based on these criteria, there are several chemicals that consistently appeared more sensitive in tadpoles for both body weight (Fig. 2a) and length (Fig. 2b). The fungicides myclobutanil and flutolanil both have unbounded LOAECs for weight and length and were >10-fold more sensitive in tadpoles. This pattern appeared in both FSTRA and ELS data (Fig. 2). Several pyrethroids (bifenthrin, cypermethrin, and cyfluthrin) had LOAECs that are also >10-fold more sensitive in tadpoles and are either unbounded in fish but not tadpoles or both fish and tadpoles; however, this pyrethroid sensitivity in tadpoles was only observed in comparison to the FSTRA, not the ELS (Fig. 2).

**Figure 2.**
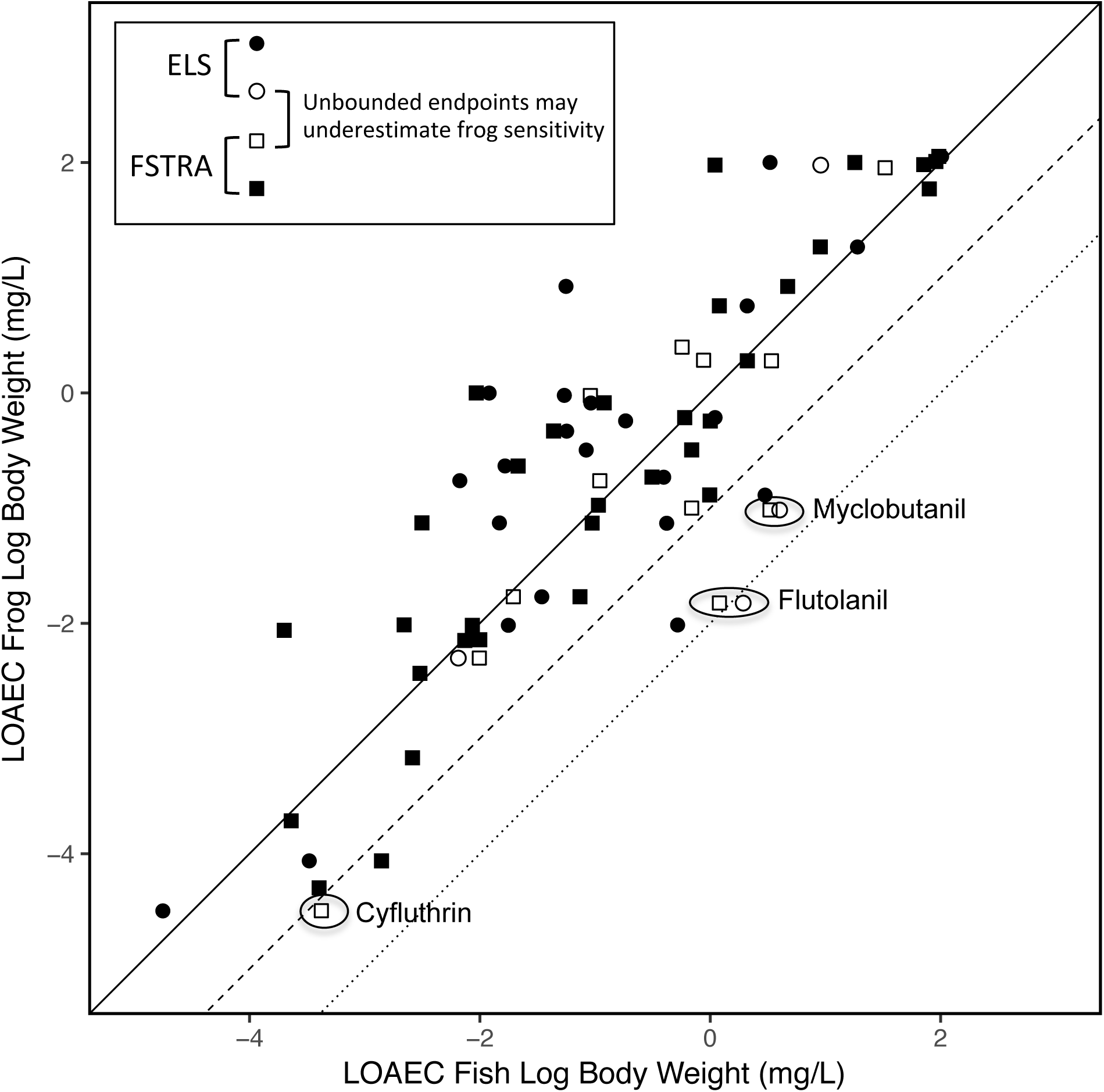
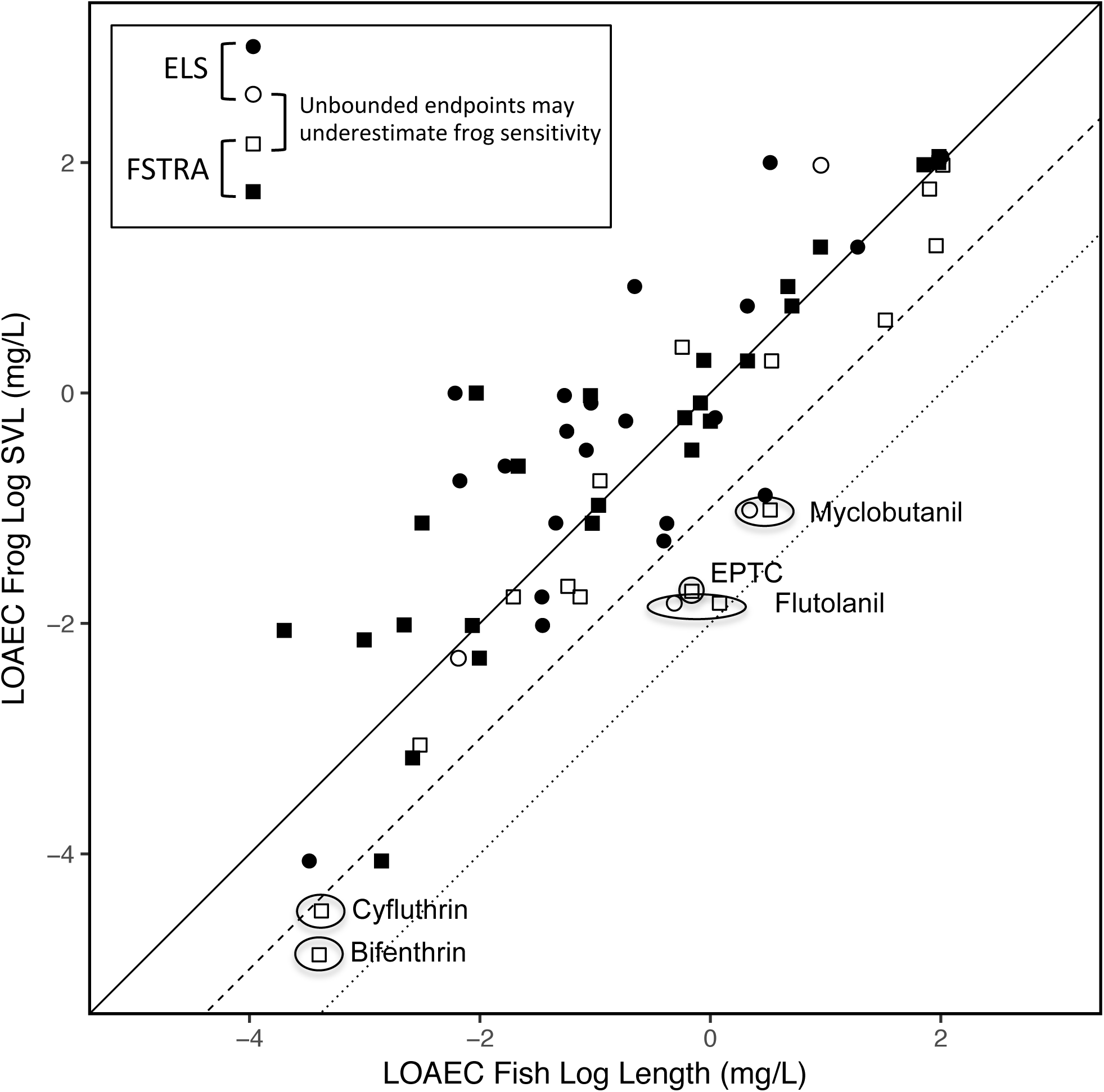
Fish-frog toxicity comparisons for (a) body weight and (b) length. Fish length corresponds to total length, while frog length corresponds to snout-vent-length (SVL). Open symbols (○/□) represent studies in which relative sensitivity is potentially underestimated in tadpoles, including cases where (1) no effects were observed at the lowest concentration in frogs, (2) no effects were observed at the highest concentration tested in fish, (3) or both. Dashed and dotted lines represent one and two order of magnitude higher sensitivity, respectively, in frogs as compared to fish.

### Comparison of FSTRA and ELS Study Designs

The linear relationship between FSTRA and ELS endpoints are slightly less robust than the relationship between FSTRA and AMA data, with *r*^*2*^ ≥0.68 (Table 2). All chemicals and growth endpoints analyzed are within two orders of magnitude between the two fish test designs (Fig. 3), except for abamectin body weight effects, which occurred at lower treatment levels in the FSTRA than in the ELS (Fig. 3a). However, there are several chemicals that were >10-fold more sensitive in the ELS and are associated with unbounded endpoints that point to possible underestimated sensitivity in the ELS: these are cases in which 1) significant effects are observed at the highest test concentration in the ELS, but not the FSTRA and/or 2) significant effects are observed at the lowest test concentration in the ELS, but not the FSTRA.

**Figure 3.**
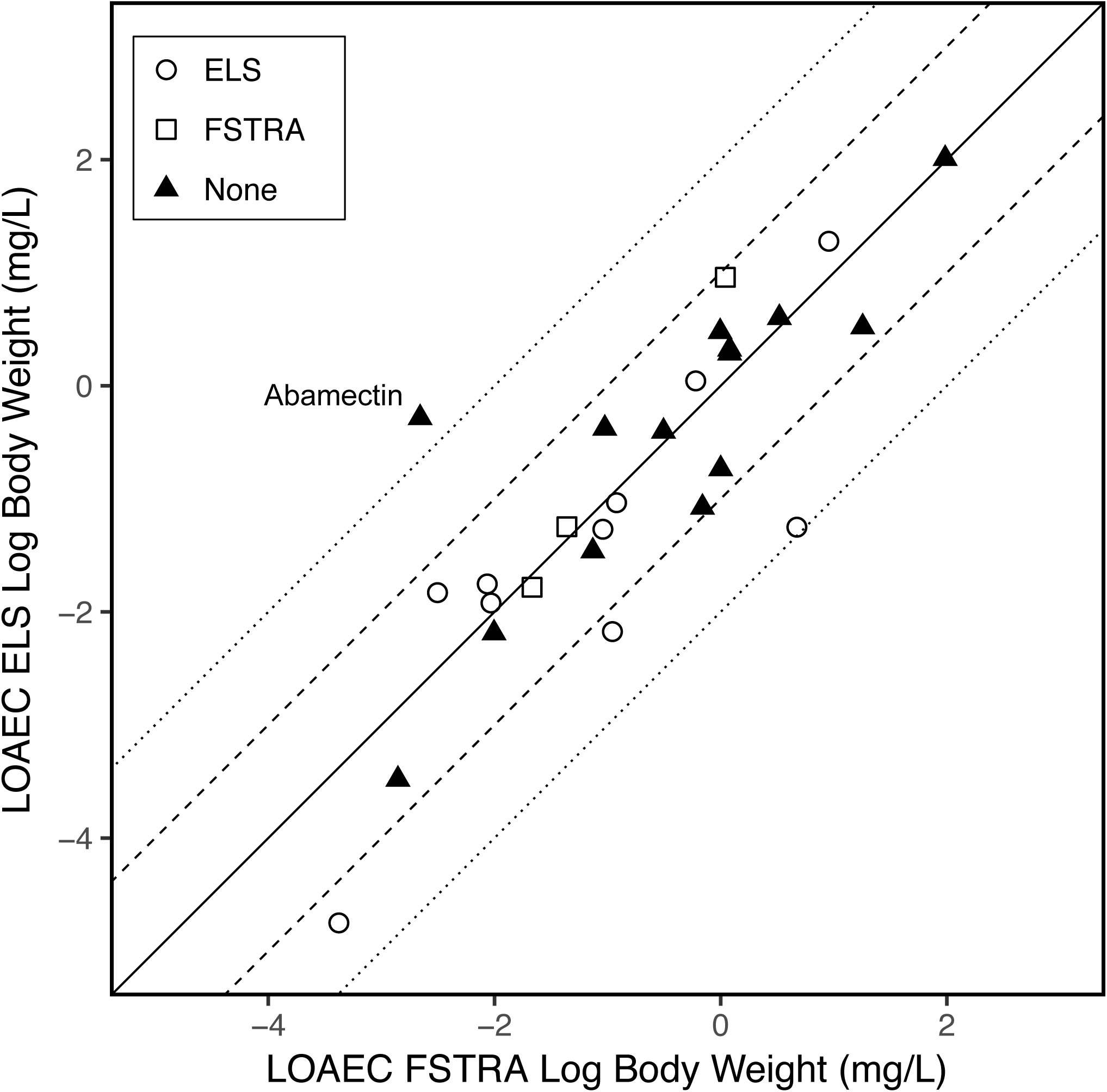
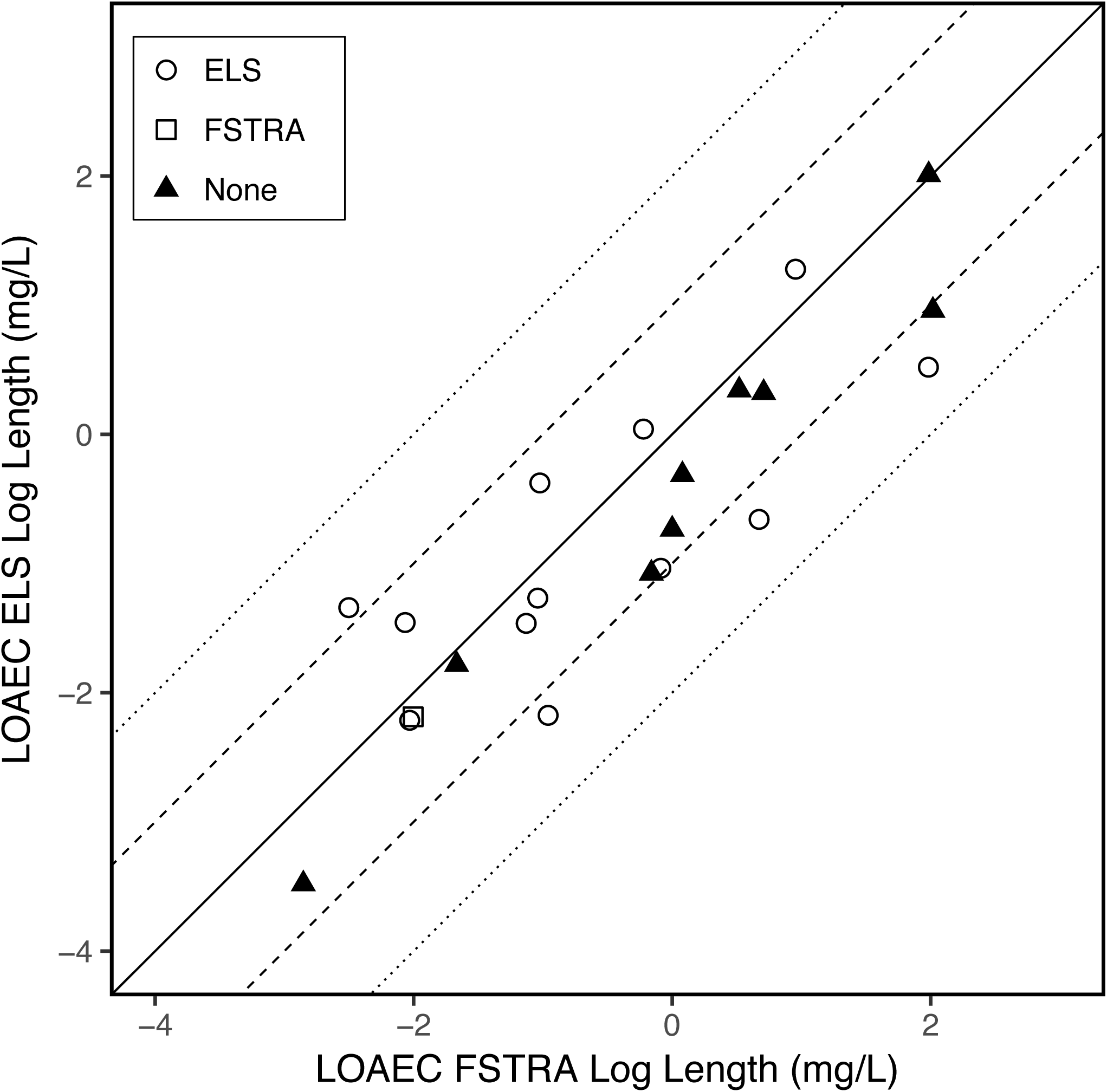
FSTRA and ELS toxicity comparisons for (a) body weight and (b) total length. Open symbols represent studies in which relative sensitivity is potentially underestimated in the ELS (○) and FSTRA studies (□), including cases where (1) no effects were observed at the lowest concentration in one of the study types, (2) no effects were observed at the highest concentration tested in the other study type, (3) or both. The closed symbol (▴) represents studies in which both ELS of FSTRA endpoints are bounded. Dashed and dotted lines represent one and two order of magnitude differences, respectively, in sensitivity between the two studies.

The t-test results indicate that paired chemical endpoint values for survival and length are significantly (p<0.05) lower in ELS studies as compared to the FSTRA (Table 2), supporting the pattern of overall greater sensitivity of the ELS study design for these endpoints, possibly due to the larger number and finer spacing of test concentrations.

## DISCUSSION

### Study Strengths and Interpretation

Our study poses significant advantages over previous datasets used to inform fish-amphibian surrogacy in ecological risk assessment. Prior studies have relied on existing published literature or public databases across which there is substantial variation in exposure method (*e.g*., static, static-renewal, flow-through), chemical concentration validation (*e.g*., nominal, measured), study duration, test species, endpoints measured, and number of concentrations tested. Moreover, there remains a dearth of consistently conducted amphibian toxicity studies over a wide range of chemicals. Here, we utilize recently generated toxicity data for pesticides from two U.S. EPA Tier I EDSP studies (FSTRA and AMA), which have similar study designs in terms of all the elements mentioned above. We also included a second, long-standing fish chronic toxicity study (*i.e.*, ELS) to provide historical grounding for our analysis. Another important attribute of all three studies examined (FSTRA, AMA, and ELS) is that each underwent at least three rounds of review by risk assessors in terms of data quality and statistical analysis. Therefore, our results provide important new information for evaluating the role of fish as surrogates for aquatic-phase amphibians in ecological risk assessment.

Overall, we found very similar sensitivities across apical endpoints based on the FSTRA and AMA study comparisons (Table 2; Fig. 2). Aquatic-phase amphibians are more sensitive than fish approximately half the time (44%-54% across endpoints) although these differences were not statistically significant. There is also a strong linear relationship between fish and frog endpoints (*r*^*2*^ ≥0.77), with slopes and intercepts close to 1 and 0, respectively. Moreover, the mean log differences in sensitivity across chemicals between the FSTRA and AMA is <0.1 and non-significant (p>0.05) for all endpoints. There is only a single instance (*i.e*., for the fungicide propiconazole) in which a chemical is >100-fold more sensitive in the AMA as compared to the FSTRA for any endpoint. Similar results are found when comparing endpoints from the long-standing ELS test to the AMA, although the correlation is slightly weaker and mean log sensitivity differences are slightly larger (but still non-significant) as compared to the FSTRA-AMA comparison (Table 2).

### Study Design Considerations

An important factor in comparing toxicity data among different test designs (*e.g.*, AMA vs. FSTRA or ELS) is the concentration regime in terms of range and spacing. Our results show that the AMA and FSTRA study concentrations are more evenly matched in terms of both concentrations tested (Fig. 1) and spacing. The lower mean and median ratio of highest to lowest concentrations in ELS studies indicate that this study is typically conducted at a narrower range of concentrations (generally <3.2x; (U.S. EPA, 2016)) than the FSTRA or AMA (10x; (U.S. EPA, 2009b, 2009c)). However, ELS studies typically use a five-concentration test design as compared to three concentrations in the FSTRA and AMA (Table 1). These features reflect the more definitive use of ELS studies for regulatory toxicology decision-making. On the other hand, the AMA and FSTRA studies are primarily used as Tier I screening studies to identify whether chemicals can impact endocrine-mediated signaling pathways and are not designed to yield precise NOAEC and LOAEC values, which explains their wider concentration spacing (*i.e*., 10x), despite having fewer test concentrations. Therefore, while the ELS likely provides greater endpoint precision, the FSTRA and AMA have much more similar study designs and provide a more consistent screening-level comparison than the ELS.

Of the three apical endpoints analyzed in this study, survival data showed the strongest relationship between the FSTRA and AMA studies (r^2^=0.85, slope=0.98). However, this should be interpreted with caution as the bulk of survival data across chemicals in both of these studies resulted in unbounded endpoints (Supplemental Table 1). This is not surprising since EDSP studies were designed to target thyroid and reproductive effects and related growth and morphological effects and to the extent possible avoid test concentrations considered likely to result in overt or systemic toxicity. Thus, concentrations tested were too low to result in significant mortality. The strong relationship found for survival likely depicts the very similar concentration regime used in both studies, and may suggest similar sensitivity for this endpoint. This similarity in concentrations tested is reasonable given that FSTRA and AMA studies for each chemical were likely conducted by the same in-house or contract laboratories utilizing the same set of range-finding information. Due to the preponderance of unbounded survival data, we evaluate the strength of our conclusions based on analysis of growth effects, which also show a strong positive relationship between the FSTRA and ELS versus the AMA.

Although the overall differences between fish and frogs were not statistically significant across chemicals, several chemicals exhibited >10-fold increased sensitivity in frogs relative to fish and were associated with unbounded endpoints potentially underestimating sensitivity in the frog; these include two pyrethroid insecticides (bifenthrin and cyfluthrin) and two fungicides (flutolanil and myclobutanil). The solvent (dimethylformamide) in the bifenthrin AMA study caused a significant (p<0.05) increase in weight and length as compared to the negative (water only) control; however, there were still significant reductions in length at all test concentrations as compared to the solvent control. Similarly, in the flutolanil AMA study, there were also clear solvent (dimethylformamide) effects on growth in control comparisons, but comparisons between treatment groups and the solvent control alone were not significant. There were no similar solvent effects in the AMA study for cyfluthrin or myclobutanil. Overall, this suggests that while AMA studies may have some issues with solvent-induced stimulatory effects, tadpoles may be more sensitive to these two groups of chemicals (pyrethroids and fungicides). This pattern should be explored further as the pesticide AMA database increases.

### Comparability of FSTRA and ELS Data for Fish

Since the ELS study has long been the standard test design for chronic effects to fish (OECD, 2013; U.S. EPA, 2016), its inclusion here provides important historical grounding for our analysis and provides an alternative fish study design for comparison with the AMA. In addition to different numbers of test concentrations, there are other important ways in which the FSTRA and ELS differ that could affect the pattern and magnitude of sensitivity. While the FSTRA is always conducted on *P. promelas*, the ELS can also be conducted with *O. mykiss* (Table 1; Supplemental Table 1), bluegill sunfish (*Lepomis macrochirus*), Atlantic silverside (*Menidia menidia*), inland silverside (*M. beryllina*), or tidewater silverside (*M. peninsulae*). Also, the FSTRA is conducted on early sexually mature fish while the ELS begins with embryos and is terminated prior to sexual maturity: therefore, these tests represent fish sensitivity at different life stages.

The relationship of growth data between the two fish studies (FSTRA and ELS) is not as strong as between FSTRA and AMA studies, with slopes closer to 1 and similar or higher coefficients of determination values between the two EDSP studies. This may indicate that the strength of the relationship among toxicity data has as much to do with study design as with the study organism, which is supported by the more similar concentration regimes in the FSTRA and AMA (Fig. 1). However, it is also possible that some of the discrepancy between FSTRA and ELS studies could be due to the different life stages tested in the two fish studies, or other sources of variability which should be further explored in the future. The analysis suggests though that study design may influence the magnitude of endpoints, which may support the strength of using the two EDSP studies for making more robust fish-frog comparisons.

### Comparison with Previous Studies

Overall, our results are consistent with those from Weltje *et al*. (2013). They compared a mixture of chronic toxicity endpoints for 52 chemicals compiled from the literature and found only two instances (*i.e*., carbaryl and dexamethasone) in which amphibians were 10-100-fold more sensitive than fish, and only a single case (sodium perchlorate) in which amphibians were >100-fold more sensitive than fish. Similarly, across the 45 pesticides compared between FSTRA and AMA studies, we found only 4, 6, and 2 chemicals that are 10-100-fold more sensitive in frogs for body weight, length, and survival, respectively, and only one instance (propiconazole for survival) in which frogs were >100-fold more sensitive than fish. There is no obvious congruence in the chemicals that came out particularly sensitive in our analysis (pyrethroids and fungicides) as compared to Weltje *et al*. (2013), who found that dexamethasone (corticosteroid), carbaryl (carbamate), and sodium perchlorate (inorganic) to be >10-fold more sensitive in aquatic-phase amphibians relative to fish. However, similar to sodium perchlorate, certain fungicides are known to act on the thyroid pathway in fish (Elskus, 2012) and mammals (Wolf *et al*., 2006), albeit secondary in some cases to hepatotoxicity. Thus, it is reasonable to expect that they might produce effects in the thyroid-sensitive AMA.

One departure of our results from Weltje *et al.* (2013) is that our fish-frog comparison data are clearly centered around the 1:1 toxicity line (*i.e*., slope of 1 and intercept of 0; Fig. 2), while chemical data in this previous study are skewed toward higher relative fish sensitivity. Moreover, where we found the tadpole to be empirically, if not statistically, more sensitive than fish 49%, 54%, and 44% of the time for body weight, length, and survival, respectively, Weltje *et al*. (2013) only found amphibians to be more sensitive 21% of the time on a chronic exposure basis. This discrepancy can be due to any number of factors including types of chemicals analyzed, study design, and species choice, but overall suggests that fish-amphibian comparisons using standardized datasets yield more similar results. Our results are generally consistent with Birge *et al.* (2000), who showed that amphibians were more sensitive than fish to organic compounds approximately half (48%) of the time, although they also reported a pattern of increased amphibian sensitivity to metals as compared to fish.

### Limitations and Uncertainties

Although our results indicate a strong relationship between fish and aquatic-phase amphibian chronic toxicity data, there are several limitations or uncertainties surrounding the dataset. As our dataset was limited to pesticides, future standardized datasets are needed to examine the consistency in fish-amphibian sensitivity patterns across chemical classes. Given the previous finding of increased amphibian sensitivity to metals (*Birge et al*., 2000) and phenolic compounds (Kerby *et al*., 2010) the generality of our conclusions among all chemicals must be further evaluated.

The Tier I EDSP toxicity data analyzed were from studies designed for endocrine screening and meant to be used in a weight-of-evidence approach along with a battery of other endocrine-related assays. The number and range of concentrations tested were not aimed at achieving definitive apical endpoints needed for assessing risk. This is best demonstrated by the fact that many endpoints were unbounded, especially for survival (Supplementary Table 1). Thus, the use of screening assays for basing ecological risk assessment conclusions should always be used with some caution (Wheeler *et al*., 2014).

Finally, the toxicity data utilized in our study only represent sensitivity to one amphibian (*X. laevis*) and two fish (*P. promelas* and *O. mykiss*) species, which presents several limitations to generalizability of the results. *Xenopus* are evolutionarily among the most basal anurans (Pyron and Wiens, 2011), and may not be representative of a vast array of frog species. Moreover, the divergence time of modern amphibians (Lissamphibia) is approximately 325 million years (Irisarri *et al*., 2017), and it is uncertain whether any single amphibian species can represent the sensitivity to chemicals across such a wide taxonomic group. Previous studies have already shown variability in chemical sensitivity among amphibians – especially among more distantly related species (Chiari *et al.*, 2015; Hammond *et al*., 2012; Kerby *et al*., 2010) – and there is evidence that *X. laevis* is among the least sensitive amphibian species to organic chemicals (Birge *et al*., 2000; Hoke and Ankley, 2005) although the data on other species are limited and the methods used in the studies varied widely. Therefore, future laboratory model development for other amphibian species should be considered.

### Implications for Ecological Risk Assessment

Overall, our finding of comparable sensitivity in fish and tadpole chronic toxicity studies supports previous studies (Birge *et al*., 2000; Weltje *et al*., 2013). Furthermore, almost no pesticides were >100-fold more sensitive in the frog relative to fish, which is in line with the E.U.’s 100-fold safety factor that is applied to fish data to account for potential higher sensitivity in amphibians. Among chemical classes (pyrethroids and fungicides) with high sensitivity in frogs, mechanistic studies may provide a useful comparison to the existing *in vivo* data, to explore whether this is explained by differences in chemical toxicity versus study issues such as solvent effects. Further development of robust and reliable mechanistic assays for the thyroid pathway may be especially useful in this regard, given the importance of thyroid-mediated processes to successful metamorphosis in amphibians. Implications of physical-chemical properties such as sorption should also be considered, which could plausibly impact differential uptake and subsequent toxicokinetics for highly sorptive molecules such as pyrethroids. Ultimately, given the known variation in sensitivity among amphibians, future studies should focus on capturing greater taxonomic diversity in amphibian chemical sensitivity in order to better represent this group of organisms in ecological risk assessment.

## ACKNOWLEDGEMENTS

This paper has been reviewed according to U.S. EPA’s journal clearance guidelines, but does not necessarily reflect the views or policies of the Agency.

